# HSF1-mediated Proteostasis Decline Links Aging and Sleep Disruption

**DOI:** 10.64898/2026.04.21.719983

**Authors:** Shintaro Yamazaki, Akhilesh B. Reddy

**Affiliations:** Department of Systems Pharmacology & Translational Therapeutics, Perelman School of Medicine, University of Pennsylvania, Philadelphia, PA 19104, USA; Institute for Translational Medicine and Therapeutics, Perelman School of Medicine, University of Pennsylvania, Philadelphia, PA 19104, USA; Chronobiology and Sleep institute (CSI), Perelman School of Medicine, University of Pennsylvania, Philadelphia, PA 19104, USA

## Abstract

Sleep disruption increases with age and is associated with adverse age-related outcomes, yet the molecular mechanisms linking these phenomena remain unclear. Here, through integrative analysis of human and mouse transcriptomic and proteomic datasets, we identify proteostasis-related pathways whose aging trajectories align with transcriptional responses to chronic sleep disruption across tissues and cell types. In the human prefrontal cortex, gene expression exhibits coherent age-associated directional shifts. Across human peripheral blood following sleep restriction and multiple aging mouse tissues and cell types, proteostasis pathways exhibit concordant downregulation. Among these, heat shock response pathways emerge as the most persistent and cross-modal signatures, with components of the heat shock factor 1 (HSF1)-mediated proteostasis network displaying diminished inducibility with age and chronic sleep insufficiency, in contrast to transient activation following short-term sleep deprivation. This attenuation is particularly pronounced in neurons, where age-associated suppression of HSF1 target programs indicates selective vulnerability of neuronal proteostasis. Spatial and single-cell analyses map this vulnerability to hippocampal circuits during aging and to superficial cortical layers and glutamatergic neurons in Alzheimer’s disease. These findings support a model in which repeated sleep disruption progressively reduces the inducible capacity of proteostatic stress responses, shifting from adaptive activation to progressive attenuation and accelerating age-related decline in proteome maintenance. Consistent with emerging functional evidence, this identifies HSF1-mediated proteostasis as an integrative axis linking sleep stability and molecular aging, suggesting a self-reinforcing relationship in which sleep disruption and proteostasis decline reciprocally exacerbate one another. These results connect transient molecular responses to sleep perturbations with long-term aging trajectories, revealing a systems-level mechanism through which cumulative sleep disruption may increase vulnerability during aging.

## Introduction

Aging is a cumulative process characterized by progressive and selective changes in cellular and physiological function and network organization ^1,2^. Sleep is a fundamental biological process that supports cellular homeostasis and organismal integrity ^3,4^. With advancing age, sleep becomes increasingly disrupted, characterized by fragmentation, reduced efficiency, and altered architecture ^5,6^. These changes are widely observed across populations and are associated with adverse health outcomes. Disrupted sleep in midlife has also been linked to increased risk of cognitive decline later in life, independent of major confounders ^7,8^. However, despite extensive epidemiological and physiological characterization, the molecular basis underlying the relationship between sleep disruption and aging remains incompletely understood.

Proteostasis, the maintenance of protein homeostasis through coordinated regulation of protein synthesis, folding, and degradation, is a central determinant of cellular resilience and declines during aging ^9^. Key proteostatic pathways, including the heat shock and unfolded protein responses, are rapidly activated following acute sleep deprivation. Experimental studies have shown that transient sleep loss induces robust transcriptional activation of heat shock proteins and other proteostasis regulators. Notably, these responses appear to be driven by sustained neuronal activity associated with extended wakefulness, independent of glucocorticoid-mediated systemic stress or temperature changes ^10,11^, indicating that proteostatic pathways are directly coupled to sleep-wake dynamics rather than reflecting a generalized stress response. Moreover, heat shock factor 1 (HSF1), a master regulator of the heat shock response, has been implicated in the regulation of sleep stability across species ^12^.

These observations support a direct link between sleep-wake dynamics and proteostatic regulation. In parallel, aging is associated with a progressive decline in proteostatic capacity across tissues ^13,14^. This decline is not uniform, as other stress-responsive pathways such as inflammatory signaling often increase with age, indicating selective remodeling rather than global functional loss. Declining proteostasis has been strongly linked to age-related neurodegenerative diseases such as Alzheimer’s and Parkinson’s disease, which themselves are frequently accompanied by sleep disruption ^15,16^. Despite this, it remains unclear how sleep-wake-coupled proteostatic responses relate to long-term age-associated decline in proteostatic capacity. Direct experimental evidence is limited, owing to the practical, ethical, and logistical constraints of sustaining controlled sleep disruption across the lifespan.

Consequently, a key unresolved question is whether chronic sleep disruption engages molecular programs that converge with aging trajectories, potentially representing a behaviorally-modifiable component of the aging process. Addressing this question requires integration across biological scales, from acute molecular responses to behavioral perturbations to gradual changes across the lifespan. Recent advances in large-scale transcriptomic and proteomic datasets across tissues, cell types, and species provide an opportunity to examine these relationships in a cross-timescale context. Here, we integrate publicly available human and mouse datasets capturing both sleep disruption and aging to examine their relationship at the level of biological pathways. We identify proteostasis-related pathways whose aging trajectories align with responses to chronic sleep disruption across multiple tissues and cell types. Notably, components of the heat shock response exhibit signatures consistent with reduced inducibility in aging, paralleling the attenuation observed with repeated sleep disruption. Furthermore, the associated molecular signatures display regional and cell-type specificity in the brain, aligning with patterns of vulnerability observed during aging and Alzheimer’s disease progression. These findings reveal a cross-timescale relationship linking transient molecular responses to sleep perturbation with long-term aging trajectories, suggesting that cumulative sleep disruption may accelerate progressive proteome maintenance decline during aging.

## Materials and methods

### Sleep-restriction transcriptome analysis

Gene expression data from Möller-Levet *et al*. ^17^ were processed to gene-level by collapsing probes to gene symbols. Expression was modeled using linear models including circadian phase and sleep condition. Differential effects of sleep restriction (SR) were estimated as log_2_ fold change with empirical Bayes moderation (limma), and genes were classified as SR-up or SR-down (adjusted P < 0.10). Gene set enrichment was performed using CAMERA (limma) with Hallmark gene sets (MSigDB). Reactome pathway enrichment was performed using over-representation analysis (clusterProfiler), with Benjamini-Hochberg correction.

### Human prefrontal cortex aging analysis

Gene expression data from adult human prefrontal cortex ^18^ were analyzed using linear models with age as the main effect and adjustment for sex, postmortem interval (PMI), RNA integrity (RIN), and batch (limma). Empirical Bayes moderation was applied, and age-associated coefficients were extracted for each gene. Genes were classified as up- or down-regulated with age based on FDR < 0.05 and a minimum effect size threshold (log_2_ change per year). For visualization, gene expression values were z-scored across samples. Age-dependent trends were visualized using loess smoothing. Gene-set trajectories were computed as the mean z-scored expression of genes within each directionality class (up-with-age or down-with-age) and plotted as a function of age

### Alignment of aging trajectories with sleep disruption

Single-cell RNA-seq data from the Tabula Muris Senis atlas ^19^ were aggregated into pseudobulk profiles by averaging gene expression within each brain cell type and age group. Aging directionality for each gene was defined as the slope from linear regression of expression versus age (expr ∼ age). Gene-by-cell-type slope matrices were visualized as heatmaps with a diverging color scale centered at zero.

For neuronal analysis, pseudobulk expression profiles were generated across ages, z-scored per gene, and visualized using loess-smoothed trajectories. Human sleep-restriction (SR) RNA-seq data were used to define gene-level directionality (log_2_ fold change). Genes overlapping between datasets were classified as aligned when aging slope and SR logFC had the same sign. Reactome pathway enrichment was performed on aligned gene sets using a hypergeometric test (clusterProfiler), with multiple testing correction by the Benjamini-Hochberg method. Enrichment results are reported as -log_10_ (FDR).

### Multi-omics aging trajectory and pathway concordance analysis

Mouse aging multi-tissue RNA-seq and proteomics datasets ^20^ were analyzed across ages (6, 15, 24, and 30 months). For each tissue and assay (RNA, whole-tissue lysate [WTL], and low-solubility fraction [LSF]), aging directionality was defined as the slope from linear regression of expression or abundance versus age. Human sleep-restriction (SR) RNA-seq data were used to define gene-level directionality (SR_up or SR_down; adjusted P < 0.10). Mouse genes were mapped to human gene symbols, and overlapping genes were retained for cross-species analysis. RNA-protein directional coupling was assessed by classifying genes as increasing or decreasing with age in each assay and quantifying concordance using Fisher’s exact test, reported as odds ratio (OR). Genes showing consistent age-related decline across RNA and protein layers within each tissue were identified and further filtered for alignment with SR directionality (aging slope < 0 and SR logFC < 0). Reactome pathway enrichment (MSigDB C2:CP:REACTOME) was performed on aligned gene sets using a hypergeometric test with Benjamini-Hochberg correction. Enrichment was quantified as -log_10_(FDR), and pathways reproducibly enriched across tissues (≥2 tissues) were retained for visualization.

### Spatial transcriptomic and cell-type-resolved vulnerability analyses

Mouse lifespan spatial transcriptomic data across the lifespan were obtained from Wu *et al*. ^21^ and analyzed across young, middle, and old age groups. Human single-nucleus RNA-seq, effect-size, and gene-dynamic matrices for middle temporal gyrus (MTG) cell populations were obtained from SEA-AD (mean_expression.h5ad, effect_sizes.h5ad, gene_dynamic_space.h5ad), together with MTG MERFISH spatial data (SEAAD_MTG_MERFISH.2024-12-11.h5ad). A brain aligned-decline gene set (SR-Aging signature) was defined from genes showing concordant downregulation across chronic sleep restriction and aging; higher-confidence analyses further required concordant decline across transcript and protein layers.

SR-Aging scores were computed at the spot level using Seurat AddModuleScore after intersecting aligned-decline genes with detected features; human gene symbols were converted to mouse-style capitalization before intersection where required. Spot-level scores were aggregated by section and age to obtain mean and median section-level scores. Global age effects were tested using the Kruskal-Wallis test, followed by pairwise Wilcoxon rank-sum tests with Benjamini-Hochberg correction. Scores were also summarized across annotated anatomical regions and broader anatomy classes. Region-level age effects were calculated as group means within each region and age, and age-associated changes were expressed as pairwise differences (middle versus young, old versus young). These regional contrasts were projected back onto representative sections for visualization of age-dependent decline.

Baseline SR-Aging enrichment was quantified in SEA-AD cell groups by z-scoring expression of detected aligned-decline genes across groups and averaging per group to obtain a proteostasis decline score. AD-associated effects were quantified by averaging gene-level effect sizes to derive a mean slope and fraction of declining genes per group. Gene-dynamic analyses computed early and late mean effect sizes, fractions of declining genes, and late-minus-early changes, with paired Wilcoxon signed-rank tests used for early-late comparisons. Baseline SR-Aging scores were integrated with AD-associated slopes across cell groups, and their association was assessed using Spearman correlation. Cell groups with highest baseline scores were defined as vulnerable supertypes and mapped onto MERFISH data using shared annotations. Spatial distributions were summarized across cortical depth, annotated layers, and sections, with layer-level fractions computed by within-layer normalization.

For spatial projection, cell-type-specific AD effect sizes were combined with local cell-type composition to generate spot-level vulnerability scores. Values were quantile-clipped for visualization, and spots were annotated by cortical layer and white matter compartments. High-vulnerability regions were defined by upper-quantile thresholds, and dominant cell-type contributions were assigned based on maximal local contribution.

### Data analysis and visualization

All analysis were performed in R (version 4.5.2) using ggplot2 (version 4.0.0), dplyr (version 1.2.0), ComplexHeatmap (version 2.26.1), limma (version 3.66.0), and clusterProfiler (version 4.18.4), Seurat (version 5.4.0), SingleCellExperiment (version 1.32.0).

## Results

### Sleep loss-specific proteostatic responses transition from induction to attenuation with chronic disruption and impact interconnected cellular programs

Acute sleep deprivation (SD) induces robust upregulation of proteostasis-related genes, including multiple components of the heat shock proteins ^10^ (Figure 1A,B). Importantly, these inductions are independent of general (glucocorticoid-mediated) systemic stress, indicating sleep loss-specific dynamics.

**Figure 1.**
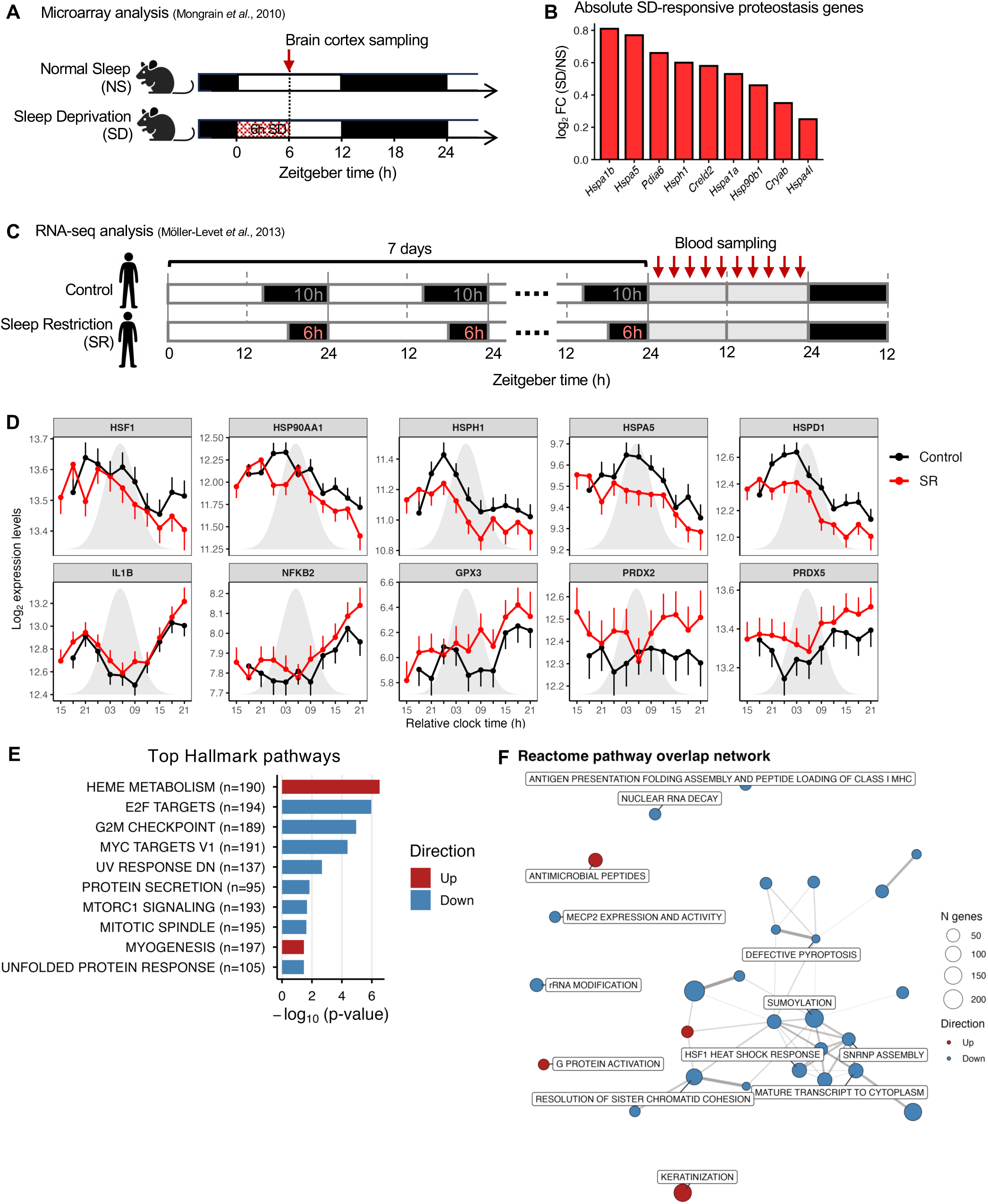
Proteostatic responses shift from induction to attenuation across increasing sleep disruption. (A) Experimental design for acute sleep deprivation (SD) in mouse cortex. Mice were maintained under a 12 h light/12 h dark cycle and subjected to SD during the early light phase, with cortical tissue collected at ZT6. Both sham-operated and adrenalectomized (ADX) mice were analyzed. ADX removes circulating glucocorticoids, allowing identification of transcriptional responses to sleep loss that are independent of systemic stress signaling. (B) Absolute SD-responsive proteostasis genes in mouse cortex. Bar plot shows log_2_ fold change (SD vs NS) for representative genes that are significantly upregulated in both sham and ADX conditions (FDR < 0.05), indicating glucocorticoid-independent induction. These include cytosolic heat shock proteins and ER proteostasis components. (C) Experimental design for repeated sleep restriction (SR) in humans. In a within-subject crossover design, participants underwent one week of control sleep (10 h per day) and sleep restriction (6 h per day), with peripheral blood collected across circadian time points under both conditions. (D) Circadian phase-aligned expression profiles of representative genes under control (black) and sleep restriction (SR, red) conditions. Expression values are plotted relative to individual melatonin timing (dim light melatonin onset, DLMO) to account for circadian phase (gray shading indicating the melatonin profile: biological night). Heat shock protein genes show reduced expression under SR across circadian phase, whereas inflammatory and oxidative stress-related genes (*IL1B*, *NFKB2*, *PRDX* family) are maintained or upregulated, indicating selective attenuation of the heat shock response rather than global suppression of stress-responsive pathways. Values represent model-estimated means ± SEM. (E) Hallmark pathway enrichment analysis of peripheral blood transcriptomes following sleep restriction. Gene set enrichment was performed using CAMERA with a linear model controlling for circadian phase. The top 10 Hallmark pathways are shown, ranked by nominal p-value. Bars represent -log_10_(p-value), with red indicating pathways upregulated in sleep restriction and blue indicating downregulated pathways. Gene set size is indicated in parentheses. (F) Reactome pathway overlap network of SR-associated transcriptional changes. Nodes represent significantly enriched pathways, with size proportional to gene count and color indicating direction of change (red, upregulated; blue, downregulated). Edges denote gene overlap between pathways. Downregulated pathways form a densely interconnected module centered on the HSF1 heat shock response, SUMOylation, and RNA-processing pathways, indicating coordinated suppression of an integrated regulatory network.

In contrast, analysis of transcriptomic changes following repeated sleep restriction (SR) in human peripheral blood ^17^ (Figure 1C) revealed reduced expression of heat shock protein genes relative to sufficient sleep group (Figure 1D), indicating a reversal of the initial induction pattern (Figure 1B,D; Supplementary File 1). Notably, inflammatory and oxidative stress-related genes were upregulated under SR conditions (Figure 1D), suggesting that attenuation of heat shock protein genes is not part of a global stress response but instead reflects a distinct regulatory process linked to sleep-wake dynamics. To assess pathway-level consequences, we performed Hallmark gene set analysis using CAMERA ^22^, which revealed coordinated downregulation of cell cycle and growth-associated pathways, including E2F targets, G2M checkpoint, and mTORC1 signaling, together with proteostasis-related pathways such as protein secretion and the unfolded protein response (Figure 1E). These results indicate coordinated suppression of biosynthetic and protein homeostasis programs under chronic sleep restriction.

To resolve the mechanistic organization underlying these changes, we next analyzed Reactome pathways and constructed a pathway overlap network based on shared gene membership (Figure 1F). Downregulated pathways converged onto a densely interconnected module comprising RNA processing, mRNA maturation, and proteostasis-related pathways, including SUMOylation, SNRNP assembly, tRNA processing, and transcript transport. These pathways exhibited extensive gene overlap and formed a central network structure. The HSF1-mediated heat shock response localized within this module alongside RNA-processing and SUMOylation pathways, consistent with known coupling between SUMO-dependent transcriptional regulation and the heat shock response ^23^. Thus, the broad program downregulation observed at the phenotype level (Figure 1E) corresponds, at finer mechanistic resolution, to suppression of an interconnected network centered on the HSF1 heat shock response, SUMOylation, and RNA-processing pathways (Figure 1F), indicating that attenuation of these core genes (Figure 1D) impacts multiple interconnected programs beyond classical proteostasis pathways.

Together, these findings reveal a transition from acute induction to attenuation across increasing durations of sleep disruption, consistent with a progressive loss of inducible proteostatic capacity that extends across multiple cellular programs.

### Human brain aging reflects cumulative and directionally organized transcriptional remodeling

To characterize aging-associated transcriptional dynamics, we analyzed gene expression profiles from the human prefrontal cortex across the adult lifespan ^18^ (Figure 2A). Gene-level analysis revealed a broad distribution of age-associated effects, with significant genes spanning a range of positive and negative age coefficients (Figure 2B), indicating heterogeneous magnitudes and directions of change across the transcriptome.

**Figure 2.**
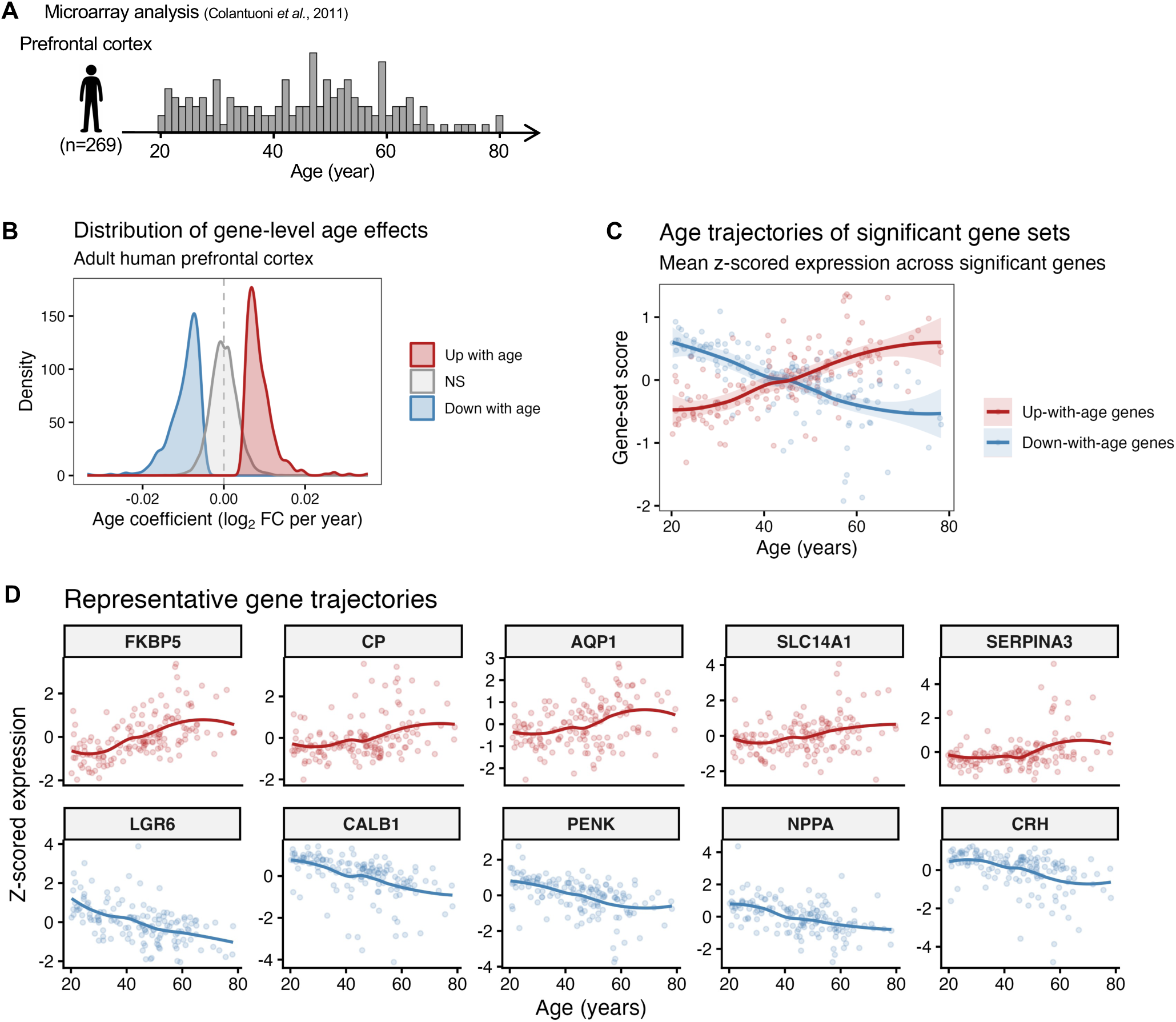
Human prefrontal cortex transcriptome exhibits directionally organized aging trajectories. (A) Study overview. Gene expression data from postmortem human prefrontal cortex were analyzed across the lifespan. The histogram shows the number of samples at each age. Samples span prenatal development through adulthood, with the present analysis restricted to adult ages. Expression values were generated using microarray profiling of dorsolateral prefrontal cortex tissue. (B) Distribution of gene-level age effects. Gene expression was modeled as a function of age, sex, PMI, RIN, and batch. Density curves show the distribution of estimated age coefficients (log_2_ fold change per year), with genes classified as up-with-age, down-with-age, or not significant. (C) Gene-set trajectories. Mean z-scored expression across genes classified as up- or down-with-age plotted against age. Each point represents an individual sample, and curves represent loess-smoothed trends with confidence intervals, illustrating opposing aggregate trajectories across the adult lifespan. (D) Representative gene trajectories. Z-scored expression of selected individual genes from the up-with-age and down-with-age groups plotted against age. Curves represent loess-smoothed trends.

Despite this variability, genes segregated into two dominant groups showing opposing trajectories across age (Figure 2C). These patterns indicate that aging is not characterized by uniform decline, but rather by coordinated shifts in distinct gene populations. Notably, these trajectories emerge progressively across the adult lifespan, with divergence apparent from early adulthood, suggesting that molecular changes underlying brain aging accumulate over time rather than arising only at later stages. Although functional changes in the aging brain are often nonlinear, with more pronounced shifts occurring in midlife ^24^, the early emergence of transcriptional shifts suggests that some molecular aging processes are initiated well before their functional consequences become apparent.

Representative gene trajectories illustrate consistent age-associated trends across individual genes (Figure 2D). Genes increasing with age, including *FKBP5*, *CP*, *AQP1*, *SLC14A1*, and *SERPINA3*, are associated with inflammatory and metabolic processes ^25–29^, whereas genes decreasing with age, such as *LGR6*, *CALB1*, *PENK*, *NPPA*, and *CRH*, are linked to regenerative, calcium handling and neuroendocrine functions ^30–34^. These opposing trajectories suggest functional partitioning between gene sets and point to molecular substrates underlying functional changes in the aging brain ^35^.

These results demonstrate that aging in the human brain is characterized by directionally organized gene expression changes with functional partitioning, reflecting cumulative molecular remodeling across the lifespan. Such organization suggests that transcriptional changes induced by persistent physiological perturbations may incrementally influence these aging trajectories in a direction-dependent manner, with concordant changes accelerating and opposing changes mitigating aspects of molecular aging.

### Aging and sleep insufficiency-induced signatures resolve to HSF1-mediated proteostasis decline as a vulnerable axis in neurons

Building on the observation that both sleep restriction (SR) and aging-associated transcriptional changes are directionally organized, we next asked whether SR engages molecular programs that align with these aging trajectories. Such concordant changes would be expected to amplify ongoing aging-associated decline, thereby identifying pathways and cell types that are particularly vulnerable to sleep loss.

To test this, we examined aging-associated pathway changes across mouse tissues and assessed their overlap with transcriptional signatures induced by SR (Figure 3A). Across tissues, aging was associated with coordinated shifts in proteostasis-related pathways, including protein secretion, unfolded protein response, and related stress-response programs (Figure 3B). These pathways overlap with those identified under SR (Figure 1E-F), indicating convergence on shared proteostatic processes.

**Figure 3.**
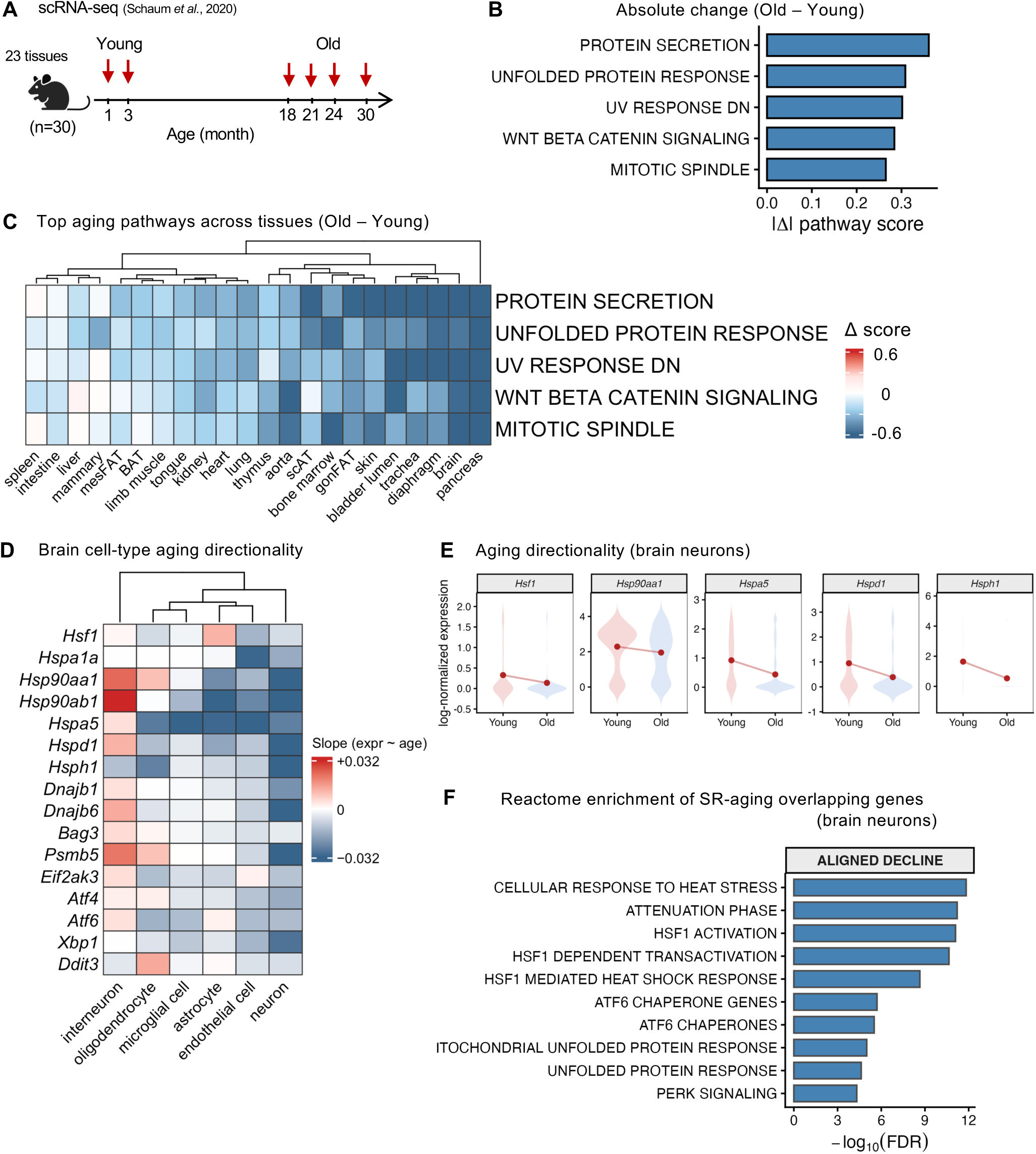
Aging and sleep restriction converge on HSF1-mediated proteostasis decline in neurons. (A) Experimental design of the aging dataset. Single-cell RNA-seq data spanning 23 mouse tissues across the lifespan (1-30 months). Young (1-3 months) and old (18-30 months) groups were defined for downstream analyses. (B) Magnitude of aging-associated pathway changes. Pathway scores were computed per tissue, and the absolute change between old and young (|Old - Young|) was used to rank pathways. Bars indicate mean absolute changes across tissues. (C) Heatmap showing the top five pathways with the largest absolute aging-associated changes (Old ≥18 months vs Young ≤3 months) across mouse tissues. Pathway activity scores were computed from tissue-level pseudobulk transcriptomes and summarized as the mean expression of genes within each pathway. Values represent Δ pathway score (Old - Young) for each tissue, computed as the difference in mean expression of pathway genes between age groups. Blue indicates decreased pathway activity with age, and red indicates increased activity. Tissues are hierarchically clustered based on pathway changes, while pathways are displayed in order of effect size. (D) Heatmap showing brain cell-type-specific aging directionality of selected proteostasis-related genes. Values represent the slope of gene expression change with age, estimated from pseudobulked single-cell RNA-seq profiles. Positive slopes (red) indicate increasing expression with age, whereas negative slopes (blue) indicate decreasing expression with age. Genes were grouped by functional class (HSP/chaperone, protein quality control/degradation, and UPR/ISR), and brain cell types were hierarchically clustered based on their gene-level aging trends. (E) Violin plots show the distribution of single-cell expression levels for selected heat shock protein (HSP) genes in brain neurons from young (3 months) and old (24 months) mice. Expression values represent log-normalized transcript abundance per cell. Lines connect group means to illustrate directional changes with age. Across all genes shown, expression is reduced in old relative to young neurons, indicating a coordinated decline in proteostasis-related transcriptional programs. (F) Reactome enrichment of SR-aging overlapping genes in brain neurons. Genes were first classified by aging direction using linear slopes of expression across age (expr ∼ age). Sleep restriction (SR) effects were defined by logFC (SR vs control) from limma with circadian phase adjustment. Genes showing consistent directionality between aging and SR (aligned decline: slope < 0 and SR logFC < 0) were selected. Reactome pathway enrichment was then performed using these overlapping genes against a background of all expressed genes. Bars represent pathway significance as -log_10_(FDR), derived from enrichment p-values corrected for multiple testing. Pathway names are simplified representations of Reactome terms.

Although these changes were observed across tissues, their magnitude varied. Tissue-resolved analysis revealed that brain exhibits consistent negative shifts in proteostasis-related pathway activity (Figure 3C), suggesting that aging-associated proteostatic decline is not uniform but may be more pronounced in specific tissues.

To further resolve this, we next assessed whether this decline is cell-type-specific. Analysis across brain cell populations showed that proteostasis-related genes generally decrease with age across cell types, with the exception of interneurons. In neurons, those genes showed a consistent downward pattern across ages, whereas non-neuronal cell types exhibited weaker or more variable changes (Figure 3D). This identifies neurons as a particularly vulnerable compartment for aging-associated proteostatic decline. Gene-level trajectories in neurons, core heat shock and chaperone-related genes, including *Hsf1*, *Hsp90aa1*, *Hspa5*, *Hspd1*, and *Hsph1*, exhibited age-associated decreases (Figure 3E), indicating that pathway-level changes are reflected at the level of those key proteostasis regulators.

Finally, genes showing age-associated decline in neurons were compared with genes altered under SR. The overlapping set was significantly enriched for HSF1-mediated proteostasis pathways, including cellular response to heat stress, HSF1 activation, attenuation phase, and related chaperone programs (Figure 3F; Supplementary File 2). These concordant changes indicate that aging and sleep disruption converge on a shared molecular axis centered on HSF1-dependent proteostasis in neurons, through which sleep behavior may selectively influence progression of aging.

### Multi-omic convergence identifies systemic proteostasis decline aligned with sleep restriction

Steady-state mRNA-protein abundance correlations are generally modest, whereas protein abundance more directly reflects functional biological output ^36,37^. Therefore, determining whether aging-associated proteostasis decline is recapitulated at the protein level is critical for interpreting transcriptomic changes in the context of biological function, particularly given evidence for post-transcriptional buffering between RNA and protein levels ^37^. We analyzed integrated bulk RNA-seq and proteomic datasets across multiple mouse tissues spanning the lifespan ^20^ (Figure 4A; Supplementary File 3). Aging-associated directionality of change (increase or decrease over time) showed significant concordance between RNA and protein layers across tissues (Figure 4B). This suggests that, over long timescales, transcriptional trends are reflected at the proteome level despite modest steady-state correlations, consistent with recent multi-omic studies highlighting the value of directional integration for identifying coordinated cross-layer signals ^38^. The magnitude of concordance was comparatively modest in the brain (odds ratio (OR) = 1.65), whereas peripheral tissues exhibited stronger coupling, with consistent propagation of transcriptional changes to both soluble and low-solubility proteome fractions.

**Figure 4.**
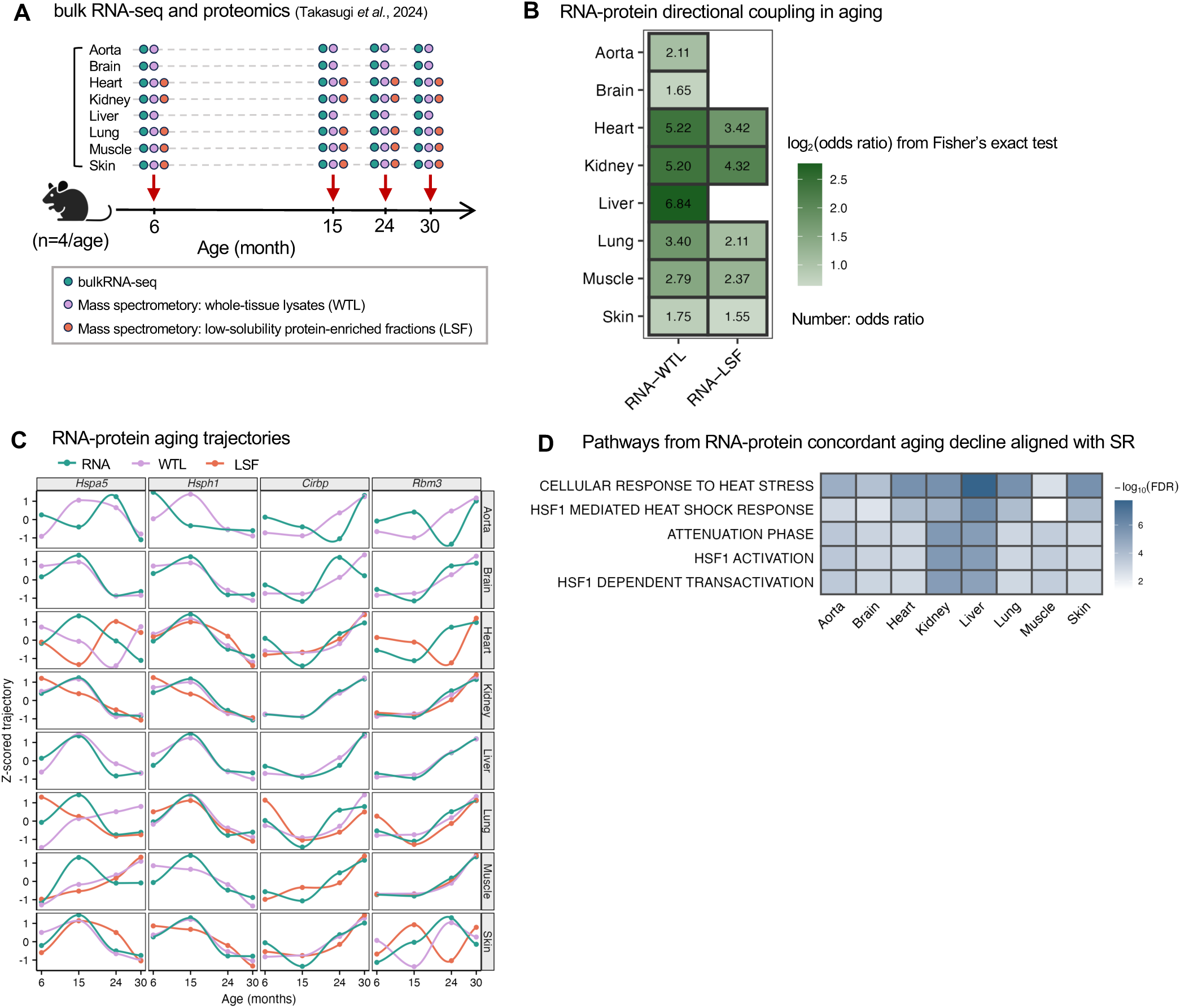
Multi-omics convergence of aging and sleep restriction reveals systemic HSF1-mediated proteostasis decline. (A) Schematic of the aging multi-omics dataset. Bulk RNA-seq and proteomics (whole-tissue lysate, WTL; low-solubility fraction, LSF) were measured across eight mouse tissues (aorta, brain, heart, kidney, liver, lung, muscle, and skin) at 6, 15, 24, and 30 months (n = 4 per age). (B) RNA-protein directional coupling in aging. For each tissue, concordance of aging direction (slope of expression vs. age) between RNA and protein layers was quantified using Fisher’s exact test. Heatmap color shows log_2_(odds ratio). Overlaid numbers denote the corresponding odds ratios, shown for significant and biologically meaningful associations (FDR < 0.05 and odds ratio ≥ 1.5). (C) Representative RNA-protein aging trajectories (6-30 months) across tissues. Z-scored expression of selected genes (*Hspa5*, *Hsph1*, *Cirbp*, *Rbm3*) is shown for RNA, WTL, and LSF. Proteostasis genes decline, whereas cold-shock genes increase, with varying RNA-protein concordance across tissues and fractions. (D) Reactome pathways enriched from genes showing concordant RNA-protein decline in aging and aligned with sleep restriction (SR). For each tissue, genes were selected based on consistent directionality across RNA and protein layers and concordant response under SR. Pathways were ranked by enrichment significance (-log_10_FDR), and recurrent pathways across tissues are shown. These include HSF1-mediated proteostasis programs, highlighting conserved molecular signatures across tissues.

Representative gene trajectories further illustrate this concordance (Figure 4C). Proteostasis-related genes, including *Hspa5* and *Hsph1*, exhibited coordinated decline across RNA and protein layers, whereas cold-shock-associated genes such as *Cirbp* and *Rbm3* showed concordant increases. These patterns suggest that aging-associated transcriptomic changes are reflected at the protein level in a subset of genes and tissues.

To identify the most robust molecular signals shared across conditions, we next focused on genes exhibiting concordant decline across RNA and protein layers and aligned with sleep restriction (SR-Aging signature). This represents a stricter criterion than transcriptome-based analyses alone, requiring agreement across molecular layers, tissues, and physiological conditions. Reactome enrichment of these concordant genes revealed consistent involvement of HSF1-mediated proteostasis pathways, including cellular response to heat stress, HSF1 activation, attenuation phase, and related chaperone programs (Figure 4D). These enrichments were observed across multiple tissues, indicating a conserved molecular signature, while remaining evident in the brain, consistent with the neuron-specific findings in Figure 3.

Collectively, these results provide multi-omic evidence that HSF1-mediated proteostasis decline represents a core axis linking aging and sleep disruption, with systemic signatures observed across tissues and corresponding biological changes evident at the protein level. Leveraging this gene set, defined as the SR-Aging signature, we next examined its behavior at higher spatial and cellular resolution.

### SR-Aging signature declines with spatial and cell-type specificity during aging and Alzheimer’s disease progression

As molecular failure precedes cellular and tissue-level dysfunction ^39^, we investigated functional relevance of SR-Aging signature expression in aging and disease contexts. Using spatial transcriptomic data from the mouse brain ^21^ (Figure 5A), we assessed regional differences in SR-Aging signature decline (Figure 5B). Early reductions were detected in the hippocampus, most prominently in CA3 and the granular cell layer (GCL) during middle age. These regions are critically involved in memory encoding and pattern separation ^40^, suggesting that early SR-Aging signature decline may contribute to initial age-associated impairments in hippocampus-dependent memory.

**Figure 5.**
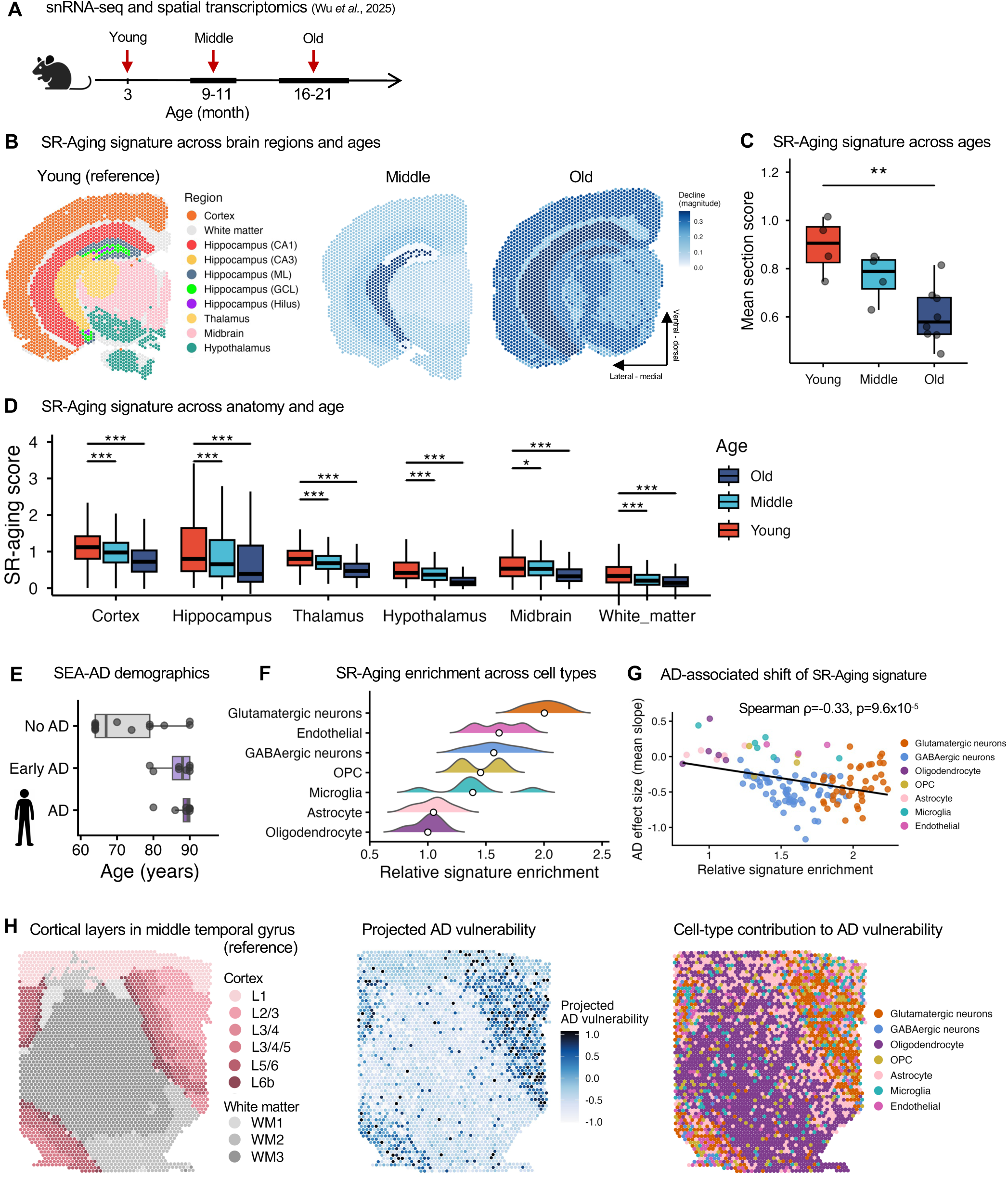
Spatial transcriptomics reveals cell-type and regional patterns of vulnerability linked to sleep disruption during aging and Alzheimer’s disease progression. (A) Overview of experimental design integrating single-nucleus RNA-seq and spatial transcriptomics across the mouse lifespan. Samples were collected at young, middle, and old ages to capture age-dependent molecular changes. (B) Spatial distribution of the SR-aging gene signature across brain sections at different ages. The SR-aging score represents the relative expression of genes jointly downregulated in sleep disruption and aging. Young brains show higher expression, while middle and old brains exhibit progressive decline. (C) Boxplots showing the distribution of mean SR-aging scores across spatial transcriptomic sections grouped by age (Young, Middle, Old). Each point represents an individual section. Boxes denote interquartile range (IQR), center lines indicate medians, and whiskers extend to 1.5× IQR. A progressive decrease in SR-aging score is observed with age. (D) SR-aging gene signature across anatomical regions and ages. Module scores are shown across major brain regions (cortex, hippocampus, thalamus, hypothalamus, midbrain, and white matter). The SR-aging signature varies by region, with strongest expression in the hippocampus and a consistent decline with age. (E) SEA-AD donor demographics. Boxplots show the distribution of donor age across diagnostic groups (No AD, Early AD, AD). Each point represents an individual donor. (F) Distribution of relative SR-aging signature enrichment across major human brain cell types. Values represent z-scored enrichment of SR-aging-associated genes within each cell type. Points indicate group means. (G) Relationship between baseline SR-aging signature enrichment and AD-associated transcriptional decline across cell groups. Each point represents a cell-type cluster. The x-axis shows baseline enrichment, and the y-axis shows AD effect size (mean slope). A negative correlation (Spearman ρ = -0.33, p = 9.6 × 10^-5^) indicates that cell types with higher baseline enrichment exhibit greater decline in Alzheimer’s disease. (H) Spatial mapping of annotated cortical layers in the middle temporal gyrus (MTG), showing laminar organization (L1-L6b) and white matter compartments (WM1-WM3). Spatial distribution of projected AD vulnerability in the MTG. Values are computed by integrating cell-type-specific AD effect sizes with local cell-type composition. Higher values indicate greater inferred susceptibility. Spatial distribution of the dominant cell-type contribution to AD vulnerability. Each location is colored by the cell type contributing most strongly to the local projected AD-associated decline.

Consistent with this, impairments in pattern separation, along with CA3 structural alterations and abnormal activity detected by MRI and fMRI, have been identified as early features of aging that precede mild cognitive impairment (MCI) ^19,41^. In older animals, the decline extended further, becoming pronounced in the dentate hilus (Hilus), molecular layer (ML), and CA1, and increasingly evident in the cortex. This broader propagation aligns with later-stage aging phenotypes, including deficits in higher-order cognitive functions mediated by cortical circuits ^42^. At the whole-brain level, SR-Aging signature expression showed a progressive decline with age (Figure 5C), while region-specific baseline expression varied in young animals and decreased across regions with statistical significance (Figure 5D).

Given that aging and sleep disruption are established risk factors for neurodegeneration ^43,44^, we next tested whether the SR-Aging signature captures vulnerability in Alzheimer’s disease (AD). Using SEA-AD human brain data (Figure 5E), we found that glutamatergic neurons exhibit substantially higher SR-Aging signature expression with approximately twofold greater than oligodendrocytes, indicating a stronger dependence on proteostasis-related programs to maintain cellular function (Figure 5F). SR-Aging signature expression declined more prominently in neuronal populations than in non-neuronal cells with AD progression (Figure 5G), in line with the preferential vulnerability and loss of excitatory neurons in AD ^44^.

Across cell types, baseline SR-Aging enrichment was inversely associated with AD-related transcriptional change (Spearman ρ = -0.33, p = 9.6 × 10^-5^), indicating that cell types with higher baseline signature expression undergo greater decline during disease progression. Projecting these cell-type-specific effects into spatial context revealed a laminar gradient of AD vulnerability in the middle temporal gyrus (MTG). Vulnerability was enriched in supragranular and mid-cortical layers (L2/3 and L3/4/5), while deep cortical layers and white matter exhibited relative resistance (Figure 5H). This pattern is consistent with known laminar susceptibility in AD, where superficial cortical layers enriched for glutamatergic neurons show early synaptic and transcriptional disruption ^45^.

These findings indicate that the SR-Aging signature captures a progression from early hippocampal vulnerability to later cortical involvement during aging, and identifies neuronal populations as selectively susceptible in AD, linking sleep disruption to aging-associated decline and disease progression.

## Discussion

Our results identify HSF1-mediated proteostasis decline as a shared molecular axis linking aging and chronic sleep disruption. Across species, tissues, and molecular layers, proteostasis pathways exhibit directionally concordant changes, transitioning from acute activation under sleep loss to attenuation with chronic disruption and aging. Molecular features of brain aging emerge progressively across the lifespan, with early changes detectable from young adulthood. This cumulative trajectory suggests that recurring physiological processes, such as sleep which occupies a substantial portion of life, may act as cumulative modulators of long-term molecular aging dynamics. These findings support the model that persistent sleep disruption accelerates age-associated decline in proteostatic capacity (Figure 6A). Consistent with this model, epidemiological studies show that chronic sleep disruption is associated with increased risk of age-related neurodegenerative diseases characterized by proteostasis impairment ^7,8^.

**Figure 6.**
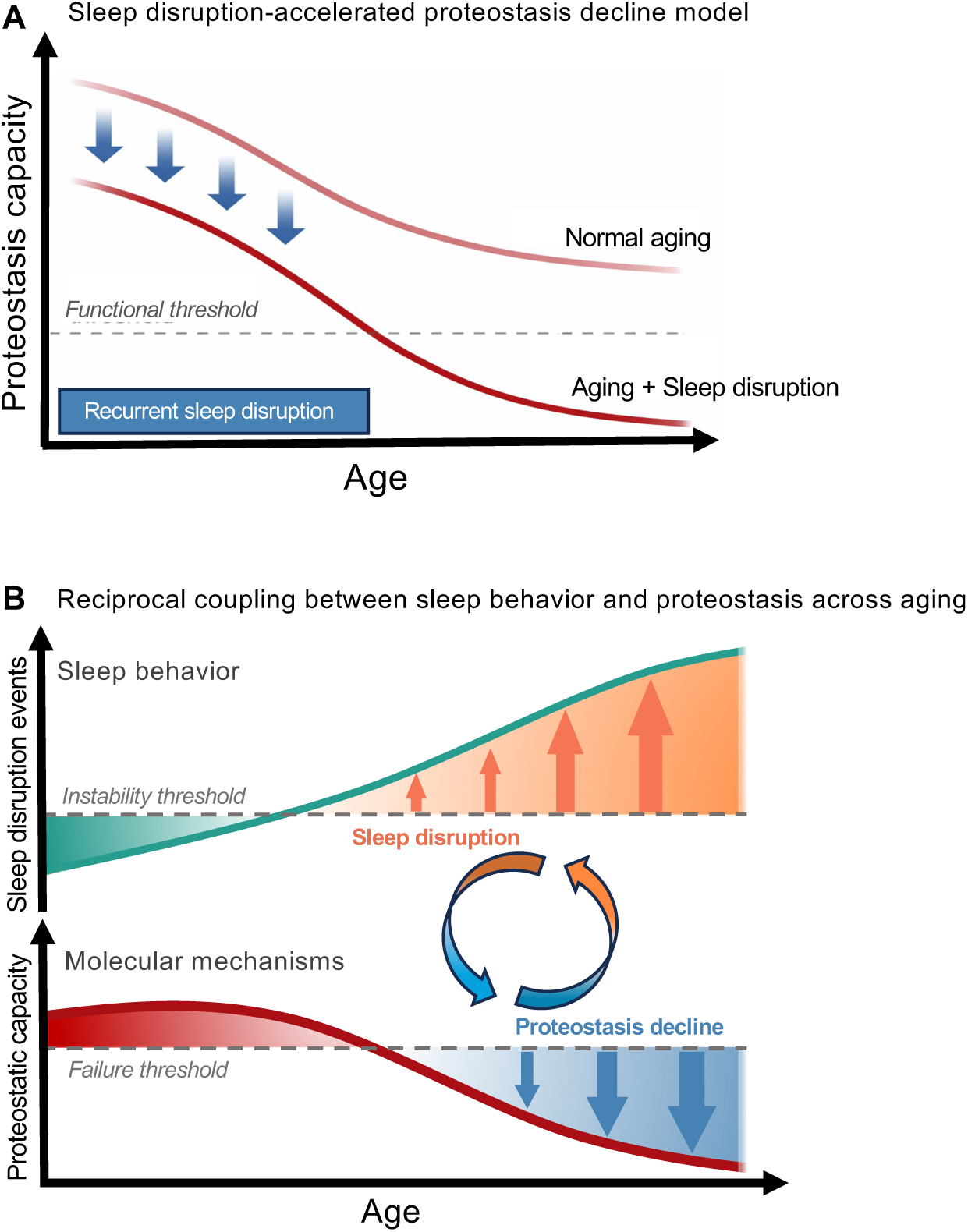
Model linking chronic sleep disruption to accelerated proteostasis decline and reciprocal sleep-proteostasis interactions. (A) Proposed effect of recurrent sleep disruption on age-associated proteostasis trajectories. Relative to normal aging, chronic sleep disruption is hypothesized to accelerate the decline in proteostatic capacity, thereby advancing the timing of functional breakdown. (B) Conceptual model of reciprocal interactions between sleep stability and proteostatic capacity. Increasing sleep instability is proposed to impair proteostasis, while declining proteostasis may further destabilize sleep, forming a self-reinforcing cycle across behavioral and molecular levels.

Notably, the core molecular signature identified here (SR-Aging signature) exhibits regional and cell-type specificity in the brain, aligning with established spatiotemporal trajectories of vulnerability ^19,41,42,45^. Given the central role of proteostasis in maintaining synaptic function, neuronal excitability, and network stability, its decline has been linked to functional deterioration in aging and neurodegenerative disease ^46–48^. The SR-Aging signature, centered on HSF1-mediated proteostasis, maps onto selectively vulnerable cell types and regions, linking molecular decline to circuit-level dysfunction. The spatial and cellular organization suggests that sleep disruption may act as a circuit-level determinant of vulnerability by progressively impairing proteostatic capacity, thereby shaping functional decline across aging and neurodegeneration.

Direct functional evidence further indicates a reciprocal interaction between sleep maintenance and proteostatic capacity via HSF1. HSF1 has been shown to modulate sleep stability, with loss of HSF1 associated with increased sleep fragmentation ^12^, indicating that proteostatic capacity can directly influence sleep-wake regulation. These observations point to a reciprocal interaction between sleep behavior and proteostasis across aging (Figure 6B). Recurrent sleep disruption attenuates proteostatic responses, while progressive decline in proteostatic capacity may in turn destabilize sleep, forming a self-reinforcing cycle. This is consistent with epidemiological and physiological observations of increasing sleep fragmentation with age ^5,49,50^, as well as evidence that inadequate sleep is associated with accelerated brain aging ^51,52^.

Brain aging is further characterized by progressive neuronal network instability ^35,53^. Given the dependence of neuronal and network function on proteostatic maintenance ^47,54,55^, reduced proteostasis provides a plausible mechanistic link between progressive sleep instability and brain aging. Moreover, studies in age-related neurodegenerative diseases, including Alzheimer’s and Parkinson’s disease models, show that sleep disruption is associated with impaired proteostasis and exacerbation of circuit dysfunction ^56,57^, further supporting a connection between sleep disruption and brain aging via proteostasis attenuation.

This cross-timescale convergence links recurrent perturbations in sleep behavior with long-term molecular trajectories within temporally ordered and spatially defined neural circuits, indicating that cumulative sleep disruption may influence the course of aging. Although causality remains to be established, HSF1-mediated proteostasis pathways represent a potential mechanistic target through which sleep behavior, as a modifiable factor, may help preserve cellular resilience during aging and disease progression.

## Acknowledgement

A.B.R. acknowledges funding from the Perelman School of Medicine, University of Pennsylvania, the Institute for Translational Medicine and Therapeutics (ITMAT) at the University of Pennsylvania. This work was supported also by NIH R35GM161590.

## Author contributions

S.Y. performed the data analysis and wrote the manuscript.

A.B.R. designed the project and analyzed results, secured funding, and wrote the paper with contributions from the other authors. All authors have reviewed the final manuscript and agree on its interpretation.

## Supplementary files

### Supplementary File 1

Gene-level differential expression and pathway enrichment results for the human sleep-restriction dataset, adjusted for circadian phase. Sheets include gene-direction statistics (Gene_direction), Hallmark pathway results and associated genes (Hallmark_results, Hallmark_genes, Hallmark_gene_summary), and Reactome pathway results and associated genes (Reactome_results, Reactome_genes, Reactome_gene_summary).

### Supplementary File 2

Reactome enrichment results for genes showing concordant directional changes between brain neuronal aging and sleep restriction. The file contains enriched Reactome terms, enrichment statistics, and contributing genes for the overlap set used to define the HSF1/proteostasis-associated decline signature.

### Supplementary File 3

Multi-omic cross-tissue results underlying Figure 4. Sheets include age-averaged whole-tissue lysate and low-solubility fraction protein values (Fig4A_WTL_mean, Fig4A_LSF_mean), RNA-protein directional coupling statistics across tissues (Fig4B_coupling_all, Fig4B_coupling_plot, Fig4B_coupling_wide), and per-tissue gene-set and Reactome enrichment summaries for concordant aging decline aligned with sleep restriction (Fig4D_gene_sets, Fig4D_enrichment, Fig4D_pathway_summary, Fig4D_heatmap_decline, Fig4D_heatmap_increase).

